# Widespread gene duplication and adaptive evolution in the RNA interference pathways of the *Drosophila obscura* group

**DOI:** 10.1101/429894

**Authors:** Danang Crysnanto, Darren J. Obbard

## Abstract

**Background:** RNA interference (RNAi) related pathways provide defense against viruses and transposable elements, and have been implicated in the suppression of meiotic drive elements. Genes in these pathways often exhibit high levels of adaptive substitution, and over longer timescales show gene duplication and loss—most likely as a consequence of their role in mediating conflict with these parasites. This is particularly striking for *Argonaute 2* (*Ago2*), which is ancestrally the key effector of antiviral RNAi in insects, but has repeatedly formed new testis-specific duplicates in the recent history of the obscura species-group of *Drosophila*.

**Results:** Here we take advantage of publicly available genomic and transcriptomic data to identify six further RNAi-pathway genes that have duplicated in this clade of *Drosophila*, and examine their evolutionary history. As seen for *Ago2*, we observe high levels of adaptive amino-acid substitution and changes in sex-biased expression in many of the paralogs. However, our phylogenetic analysis suggests that co-duplications of the RNAi machinery were not synchronous, and our expression analysis fails to identify consistent male-specific expression.

**Conclusions:** These results confirm that RNAi genes, including genes of the antiviral and piRNA pathways, have undergone multiple independent duplications and that their history has been particularly labile within the obscura group. However, they also suggest that the selective pressures driving these changes have not been consistent, implying that more than one selective agent may be responsible.

## Introduction

Gene duplication is an important process in molecular evolution, providing raw genetic material for evolutionary innovation. The evolutionary dynamics following gene duplication are often described in terms of two alternative models, ‘neofunctionalization’ and ‘sub-functionalization’ [1]. Under neofunctionalization, the functional redundancy following duplication provides relaxed selective constraint, and allows new mutations to accumulate through genetic drift. Most such mutations will reduce the functionality of the gene (resulting in pseudogenization), but some paralogs can be selected for novel or derived functions. Under sub-functionalization, the duplicates independently accumulate mutations that allow them to specialise in a subset of ancestral functions of a pleiotropic gene. Neo-functionalization leads to asymmetrical evolutionary rates among paralogs (with faster evolution in paralogs that gain derived function), whereas equal rates are expected for the latter [2]. It has been suggested that both processes have played an important role in the rapid evolution of RNA interference-related pathways, including the long- and short-term evolutionary history of the Argonautes, the effectors of RNAi [3–5].

The RNAi-related pathways comprise a range of small-RNA mechanisms best known for their roles in mediating the control of gene expression, antiviral responses, and defence against mobile genetic elements. Respectively, these include the miRNA pathway (Dicer-1 and Argonaute-1 in insects [6]), the siRNA pathway (Dcr-2 and Argounate 2 in insects [7]), and the piRNA pathway (piwi-family Argonaute AGO3 and Piwi/Aub in insects [8, 9]). In addition, RNAi-related pathways have been implicated in a variety of other biological processes, such as the control of dosage compensation [10–12] and the suppression of genetic drive [13–18]. Several genes involved in the defensive piRNA and siRNA pathways, but not genes of the miRNA pathway, display elevated rates of adaptive protein evolution. This is best studied in *Drosophila* [19–21], but is also detectable in other invertebrates [22]. It has been hypothesized that this may be a consequence of parasite-mediated ‘arms-race’ coevolution [20, 23], either through conflict with parasite-encoded immune suppressors—as widely seen in RNA viruses [24]—or in the case of the piRNA pathway, through selection for ‘re-tuning’ suppression mechanisms [25].

Adaptive evolution of RNAi pathways is partly reflected in the gain, loss, and functional divergence of Argonaute-family duplications [26]. For example, within the Drosophilidae—an important model for RNAi-related pathways of animals—Piwi has been duplicated in the lineages leading to *Phortica variegata* and *Scaptodrosophila deflexa* [3], and *Ago2* has been duplicated in those leading to *S. deflexa*, *D. willistoni*, *D. melanogaster* (where only one paralog remains – the canonical *Ago2*) and *D. pseudoobscura* [4]. This is particularly striking in the obscura group of *Drosophila* species [27], which has experienced at least 6 independent duplications of *Ago2* over the last 20 million years, with all but one of the resulting duplicates becoming testis-specific, and most displaying evidence of recent and/or ongoing positive selection [4].

Recently it has been noted that several accessory components of the siRNA and piRNA pathways have also been duplicated in *D. pseudoobscura*, including *armitage*, *asterix*, *cutoff*, *maelstrom*, *tejas* and *vreteno* [22]. In *D. melanogaster*, these proteins are engaged in a number of roles in the piRNA pathway (**Table 1**). Here we use publicly available data to reconfirm the history and expression of *Ago2* in the obscura group, and to test whether duplications in the other genes also show male-specific expression, whether duplications are contemporaneous with those of *Ago2*, and whether they too show strong signatures of adaptive protein evolution. We find no clear pattern of these duplications being coincident with *Ago2* duplications, but both *asterix* and *cutoff* duplications display increased sexual dimorphism relative to their ancestral copies, through decreased female expression. In addition, several of the gene duplicates show evidence of adaptive protein evolution in *D. pseudoobscura*, including both copies of *cutoff*, the ancestral copy of *asterix*, and the new duplicates of *tejas*, *maelstrom* and *vreteno*.

**Table 1.** **The details of RNAi accessory genes duplicated in the obscura group as reported by** Palmer *et al.* [22] *The tissue gene expression in Dmel is based on the FlyAtlas2 [28]. We report tissues with enrichment > 0.4.

| Gene | Involvement in piRNA pathway | Function of the protein product | Tissue expression in *Dmel* | Reference |
| --- | --- | --- | --- | --- |
| *armitage*<br>(*armi*) | piRNA biogenesis | A RNA helicase which unwinds the piRNA intermediates before loading into Piwi | ubiquitously expressed | [29]<br>[30] |
| *asterix*<br>(*arx*) | TGS<br>(Transcriptional Gene Silencing) | A zinc-finger protein which directly interacts with Piwi to scan and identify the transposon transcriptions as target for histone modifications | ubiquitously expressed | [31]<br>[32] |
| *cutoff*<br>(*cuff*) | piRNA transcription | Forms a complex with Rhino-Deadlock-Cutoff (Rhi-Del-Cuff) to protect uncapped non-canonical (dual-strand cluster) piRNA transcript from degradation, splicing and transcription termination | testis | [33] |
| *maelstrom*<br>(*mael*) | TGS | Act downstream of Piwi to establish histone modification and prevent the spreading of the silencing marker to the surrounding genes | brain, testis, adult female salivary gland, ovary | [34] |
| *tejas*<br>(*tej*) | Secondary piRNA production<br>(Ping-pong cycle) | A Tudor-domain protein which physically interact with *Vas*, *Spn-E* and *Aub* for a proper ping-pong cycle in the *nuage* | testis, accessory glands, adult female salivary gland, ovary | [35] |
| *vreteno*<br>(*vret*) | piRNA biogenesis | A Tudor-domain protein which essential for an early primary piRNA processing | larval brain, adult female salivary gland, ovary, testis | [36] |

## Results

### Obscura group ***Argonaute 2*** genes are duplicated and show male-biased expression

The obscura group has experienced multiple duplications of *Ago2* and it has previously been shown that these are associated with positive selection and testis-specific expression [4]. Here we reanalyzed the expression patterns and evolutionary history of these genes using publicly available RNAseq and genomic data, additionally including newly available genomic sequences from *D. algonquin* [37], *D. athabasca* [37], *D. bifasciata* and *D. miranda* [38].

In contrast to the previous qPCR analysis that failed to identify substantial or easily detectable expression of the ancestral copy in *D. pseudoobscura* (*Ago2d* [4]), we found the expression of all *Ago2* homologs in *D. pseudoobscura* was detectable at a high level in RNAseq data, and that all show significant male-bias (**Figure 1**). The *Ago2d* expression detected here is unlikely to be an artefact of cross-mapping between paralogs as we observed the reads that mapped uniquely across the gene. The male bias was largest for *Ago2e*, where expression in males is approximately 1000-fold higher than females (pMCMC<0.001; **Figure 1**), and smallest in *Ago2c* and the ancestral copy *Ago2d*, consistent with the *ca.* two-fold enrichment of the single copy of *Ago2* in male *D. melanogaster*. We also confirmed that *D. miranda*, a close relative of *D. pseudoobscura* that has not previously been analyzed, displayed a qualitatively similar pattern among those paralogs represented (**Figure 1**). In *obscura*, we found the ancestral copy (*Ago2a*) again showed slightly, but marginally significant higher expression in males (pMCMC=0.014), but that other *Ago2* proteins showed a strong male biased expression, with the largest effect for *Ago2f*, where male expression was 2000-fold higher (**Figure 1**; pMCMC<0.001).

**Figure 1.**
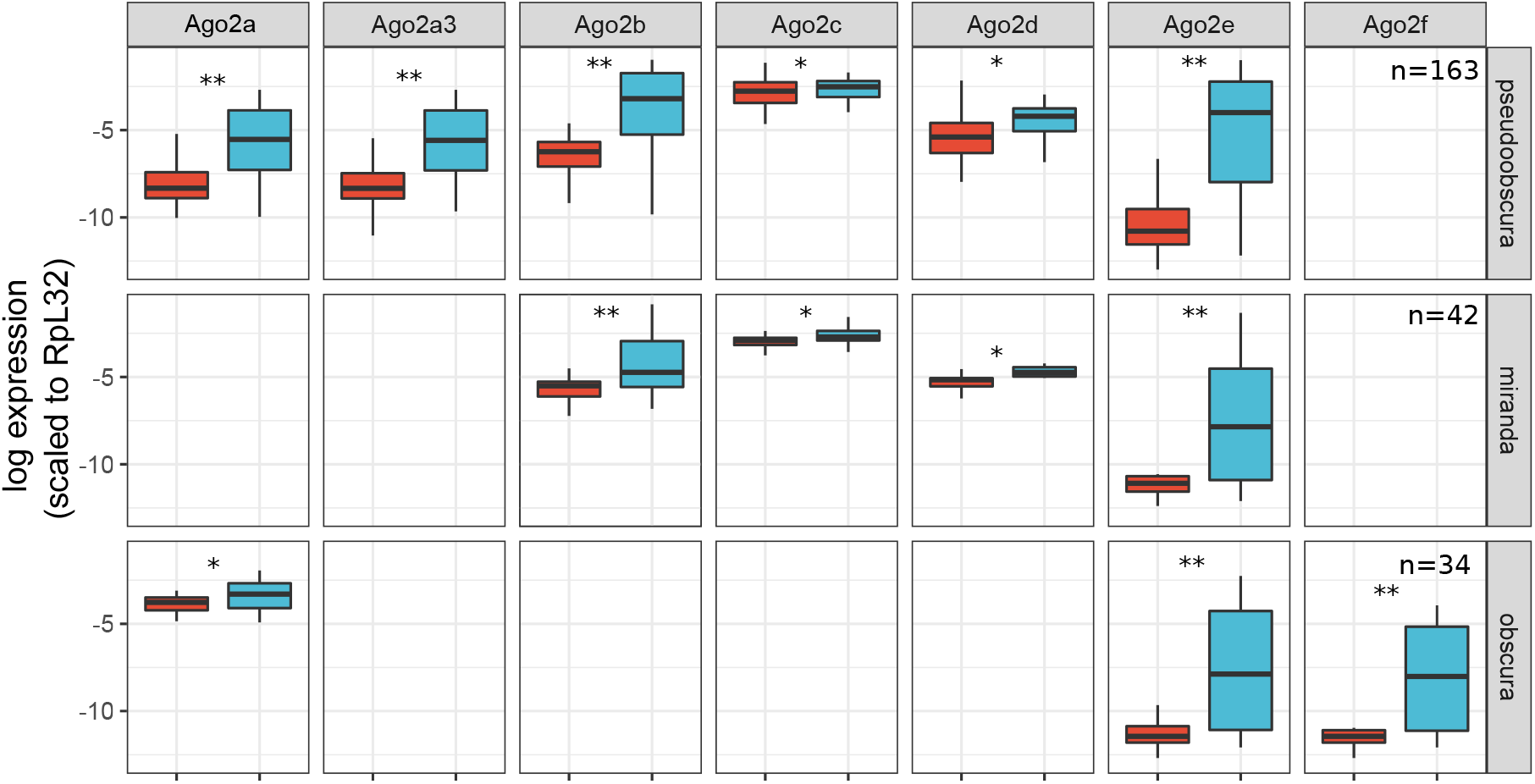
Expression profile of *Argonaute 2*. Plots show the difference in expression between female (red) and male (blue) flies, based on public RNAseq data, normalized to rpL32 and plotted on a natural log scale. The significance of differences between the sexes was assessed using a linear model fitted with MCMCglmm and is denoted by asterisks: * 0.001 < pMCMC < 0.05; ** pMCMC <= 0.001). Sample size (n) represents the number of RNAseq datasets used (combined across tissues). *Ago2d* is the ancestral copy in Dpse and Dmir, *Ago2a* is the ancestral copy in *D. obscura* and *Ago2a* is recently duplicated in Dpse become *Ago2a1* (*Ago2a*) and *Ago2a3*.

### Six piRNA pathway genes are duplicated, and ***asterix*** and ***cutoff*** duplicates show increased male-bias in their expression

Palmer *et al.* [22] recently identified six accessory piRNA pathway genes that have also experienced duplication in the obscura group (**Additional file 1 Table S1**). We could locate the duplicates for all genes in *D. pseudoobscura*, except for *armitage* where we instead identified a duplicate in the affinis subgroup but not in the pseudoobscura, obscura or subobscura subgroups. At the time of analysis, assembled genomic resources were not available for the obscura and subobscura subgroups, and our analysis of transcriptomic data did not identify duplications in those genes. Following submission, draft genome assemblies became available for *D. obscura* [39] and *D. subobscura* [40]. A search of these genomes confirmed that duplicates are undetectable for all genes except *vreteno*, but that two copies of *vreteno* are only detectable in *D. obscura* (not shown).

Where new chromosomal locations of the duplications could be determined by synteny in *D. pseudoobscura*, we found that *cutoff*, *maelstrom* and *vreteno* were duplicated from an autosome to the X chromosome, *asterix* duplicated from the X chromosome to an autosome, and *tejas* duplicated between autosomal locations. Two duplicates (*asterix* and *tejas*) lack introns, suggesting they are retro-transcribed copies created through an mRNA intermediate.

Using public RNAseq data from *D. pseudoobscura* and *D. miranda*, we found that all of the gene duplicates were expressed (**Figure 2**). *Armitage*, which was not duplicated within the newly examined lineages for which RNAseq data were available, did not show strong sex-biased expression. Similarly, *maelstrom*, *tejas*, and *vreteno* were not strongly differentially expressed between the sexes, and nor were their duplicates in *D. pseudoobscura* and *D. miranda*. In contrast, both *asterix* and *cutoff* duplicates displayed substantially reduced expression in females and slightly increased expression in males relative to the ancestral copy (**Figure 2**). For example, as previously reported from qPCR analysis [41] the paralog of *asterix* in *D. pseudoobscura* displays *ca.* 1000-fold higher expression in males than females. In both *D. pseudoobscura* and *D. miranda* those genes with overall strongly increased male-biased expression (*Argonaute-2*, *asterix*, *cutoff*, and their paralogs) had the highest expression in testis, and had reduced expression in ovaries (**Additional file 2 Figure S1**).

**Figure 2.**
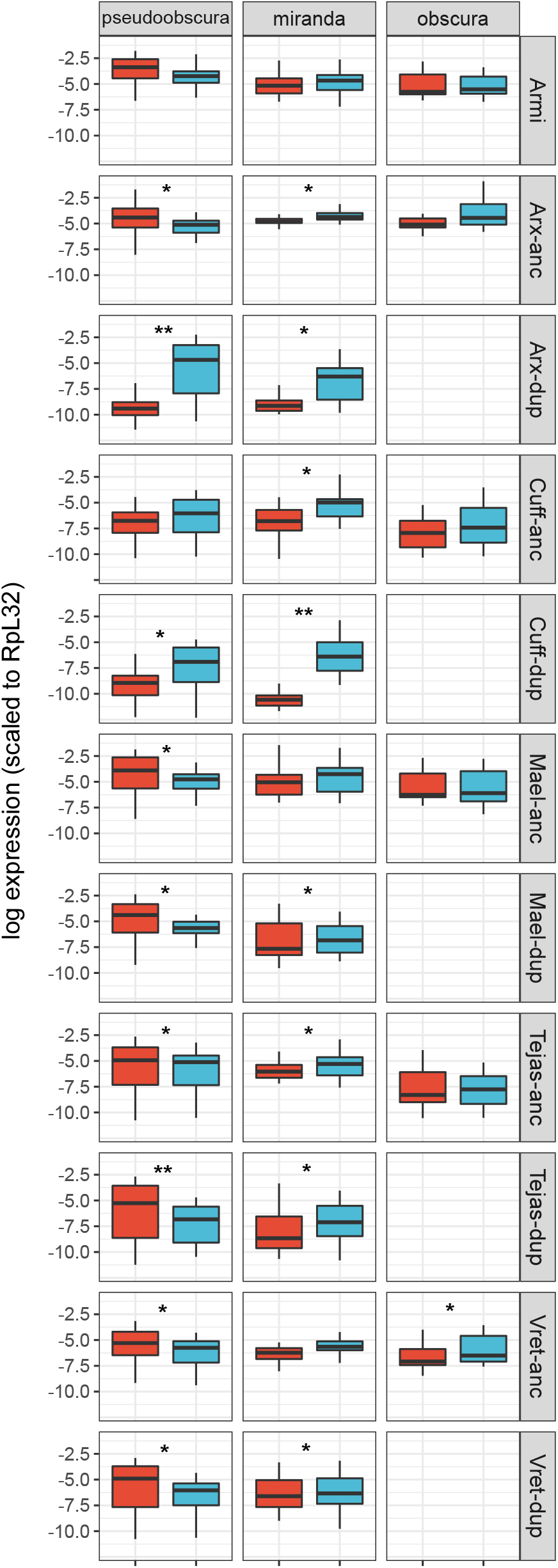
Expression profile of RNAi-accessory protein genes. Plots show the expression pattern between sex (male: blue, female:red) for genes other than Ago2; ‘anc’ ancestral copy, ‘dup’ duplicate copy as inferred by synteny. The y-axis is the natural log of normalized expression. The significance between sexes is denoted by * (0.001 < pMCMC < 0.05) and ** (pMCMC < 0.001). Sample sizes are the same as **Figure 1.** Note that *armitage, asterix, cutoff, maelstrom, tejas and vreteno* are not duplicated in the obscura subgroup.

### Adaptive amino-acid substitutions are generally more common in the duplicates

Using population genetic data from *D. pseudoobscura*, with *D. miranda a*s an outgroup, we used the McDonald-Kreitman framework and a maximum-likelihood extension to estimate the rate of adaptive substitution in protein sequences, and to test whether this rate differed between the ancestral and duplicated copies [42, 43]. Treating genes individually, we found evidence for positive selection acting on at least one paralog for each of the genes except *asterix* and *Ago2c* (p < 0.05; **Additional file 3 Table S2**). Among the ancestral copies, only *cutoff* displayed evidence of positive selection. We then tested whether the paralogs generally showed a different pattern of selection to the ancestral copies by dividing the genes into two classes (6 ancestral copies and 8 paralogs) and comparing the likelihood of models that allowed the classes to differ in the adaptive rate α (**Table 2**) [43]. The best-supported model allowed α to differ between ancestral and duplicate copies (Akaike weight: 0.81), and the second-best supported model was that in which ancestral copies experienced no ongoing positive selection (i.e. α=0; Akaike weight: 0.19), providing overall evidence that the paralogs have experienced more adaptive protein evolution. In the best-supported model, the α value was estimated to be 0.68 for the duplicate group, which more than three times larger than, the α value of the ancestral group (0.20). In case segregating weakly-deleterious variants had led to a downward bias in estimates of α, we repeated this analysis excluding all alleles with a minor allele frequency <0.125 [44], although this reduced power to the extent that few genes remained individually significant (**Additional file 5 Table S3**). We also repeated the analysis with a larger dataset PRJNA326536 [45] (**Additional file 5 Table S3**), and obtained qualitatively similar results (R^2^=0.946 for α estimates between the analyses; the second dataset, while larger, is less suitable for analysis as only the third chromosome is a direct sample from a wild population).

**Table 2.** Joint estimates of adaptive evolution across genes. Maximum-likelihood extension of MK test model fitted with different constraints on α [43]. LnL is the log likelihood of the model, AIC is the Akaike Information Criterion with corresponding relative probability as Akaike Weight (*wi*). The most supported model is in bold

| Model | Model description | LnL | AICc | Akaike weight ($w_i$) | Maximum likelihood estimate of $\alpha$ | |
| --- | --- | --- | --- | --- | --- | --- |
|  |  |  |  |  | ancestral | duplicate |
| M0 | $\alpha\text{-anc}=0, \alpha\text{-dup}=0$ | -300.61 | 633.2207 | $4.00 \times 10^{-12}$ | 0 | |
| M1 | $\alpha\text{-anc} > 0, \alpha\text{-dup} > 0$<br>$\alpha\text{-anc} = \alpha\text{-dup}$ | -284.105 | 602.2092 | $2.6 \times 10^{-5}$ | 0.539 | |
| M2 | $\alpha\text{-anc} > 0, \alpha\text{-dup} = 0$ | -301.857 | 637.7133 | $5.10 \times 10^{-13}$ | 0.047 | 0 |
| M3 | $\alpha\text{-anc} = 0, \alpha\text{-dup} > 0$ | -275.249 | 584.4975 | 0.186 | 0 | 0.618 |
| **M4** | **$\alpha\text{-anc} > 0, \alpha\text{-dup} > 0$**<br>**$\alpha\text{-anc} \neq \alpha\text{-dup}$** | **-272.774** | **581.5471** | **0.813** | **0.2** | **0.676** |

### Gene duplications were unlikely to be contemporaneous

Given the multiple duplications of *Ago2* and the piRNA pathway components in the obscura group, we hypothesized that some duplications may have occurred near-simultaneously, duplicating whole components of a pathway together. We therefore used relaxed-clock phylogenetic methods to estimate the relative timings of each duplication. In agreement with the previous analysis of Ago2 [4], we found that the duplications giving rise to *Ago2e* and *Ago2f* predated the split between the obscura and pseudoobscura subgroups, with a subsequent loss of *Ago2f* from the pseudoobscura subgroup (**Figure 3**). In contrast, we found that duplications in five of the other six genes unambiguously occurred after the obscura/pseudoobscura split, with the timing of duplication in *maelstrom* being uncertain. Briefly, *armitage* displayed a single duplication shared by members of the affinis subgroup, *asterix* and *tejas* a single duplication each in the lineage leading to *D. pseudoobscura* (which were subsequently lost in the affinis subgroup), *cutoff* a single duplication recently in the pseudoobscura subgroup, and *vreteno* a single duplication at the base of the obscura group (**Figure 4**). For *maelstrom*, the maximum clade credibility tree suggests duplication occurred very slightly prior to this split, followed by subsequent loss of one paralog the obscura subgroup (**Figure 4**). However, this was poorly supported, and a duplication that post-dates the split between the affinis and pseudoobscura subgroups, and so does not require a hypothetical loss of one paralog from the affinis subgroup, may be more parsimonious. We used the posterior distributions of split times, relative to the divergence time of the obscura and pseudoobscura subgroups, to infer whether or not duplications occurred at approximately the same time (**Figure 5**). Although the small amount of information available from single genes made relative timings highly uncertain, it is clear that few of the *Ago2* duplications could have been concurrent with the piRNA-pathway duplications (**Additional file 4 Figure S2**). However, the recent and rapid duplications within the piRNA pathway could have been concurrent, with *vreteno*, *tejas, maelstrom* and *asterix* not differing significantly, all having duplicated very close to the split between *D. obscura* and *D. pseudoobscura* (posterior overlap >0.1 in each case; **Additional file 4 Figure S2**).

**Figure 3.**
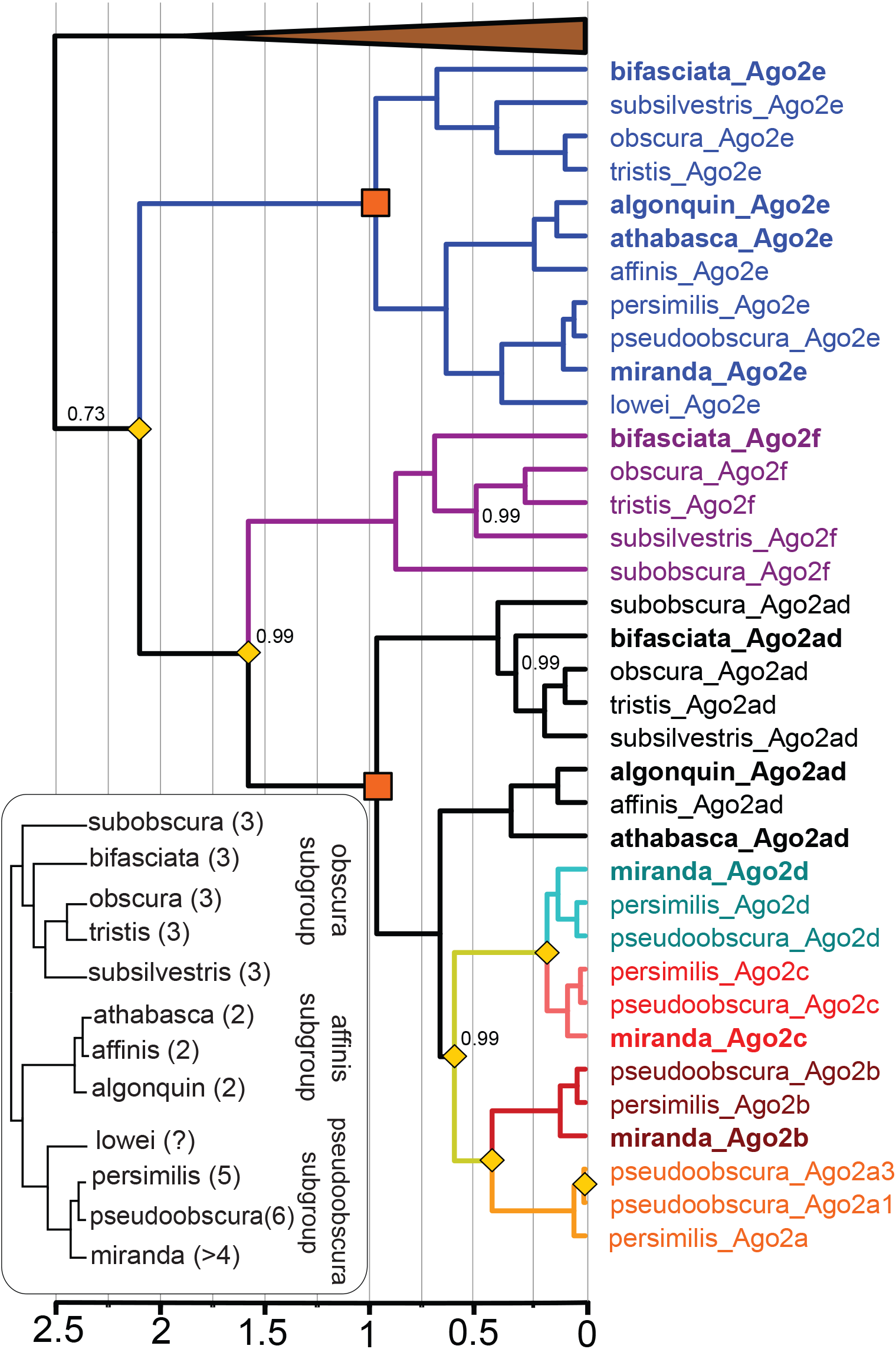
Bayesian relaxed clock gene tree for *Argonaute 2*. Duplication events are marked by yellow diamonds, species other than the obscura group are collapsed (brown triangle), and paralog clades are colored. Bayesian posterior supports are only shown for the nodes with support less than 1. Genes not previously included in the analysis of [4] are marked in bold. Time is expressed relative to the split between the obscura and subobscura subgroups (orange boxes), which was constrained to be 1 using a strongly informative prior. For reference, the obscura-group species tree inferred from concatenated ancestral copies of each gene (consistent with published phylogenies of the group [46]) is shown inset, with the inferred number of Ago2 paralogs marked next to each species.

**Figure 4.**
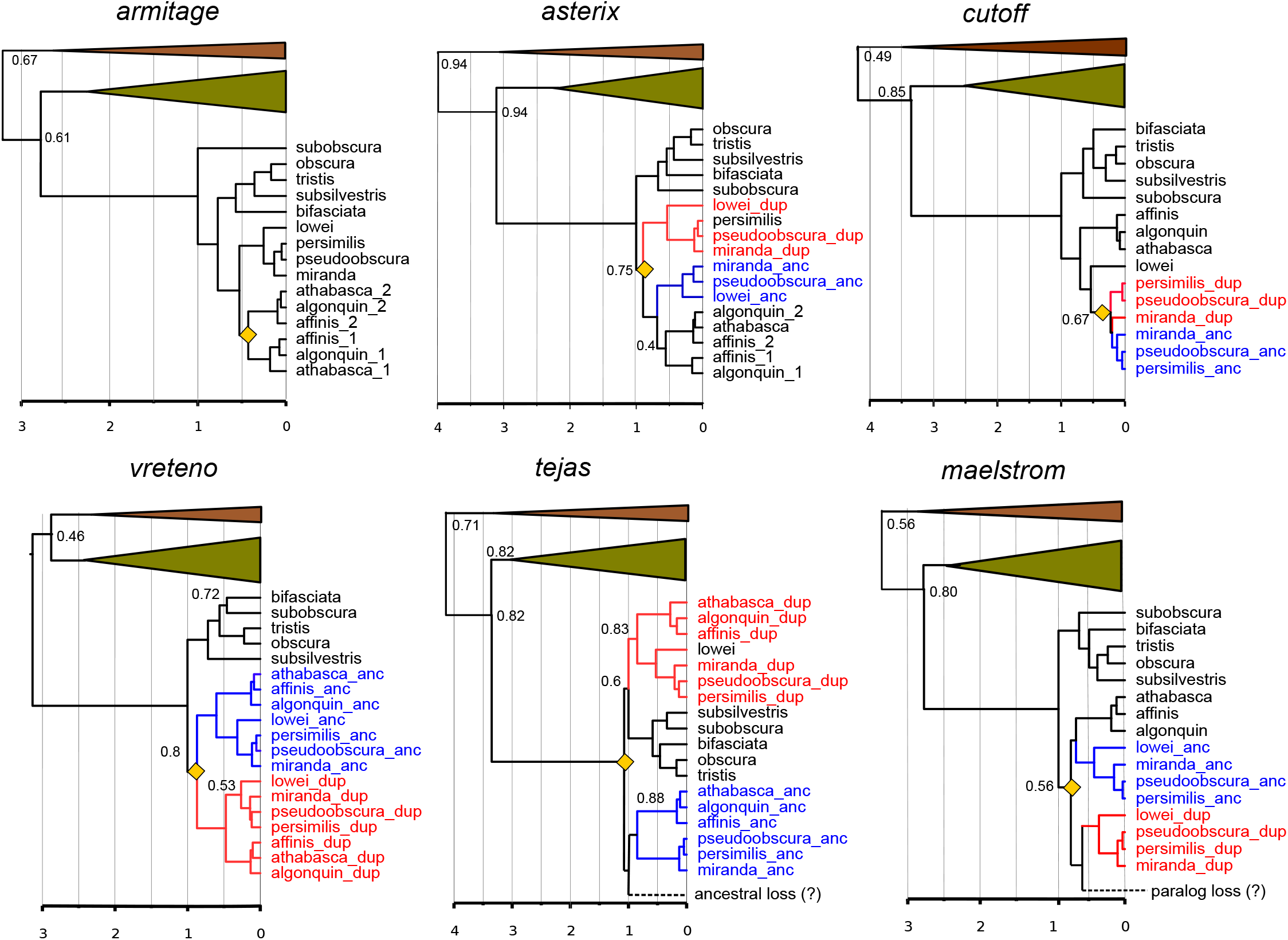
Bayesian relaxed clock trees for 6 RNAi accessory protein genes. Ancestral genes are marked by bold blue, duplicates in bold red. Yellow diamonds indicate duplication events. Species other than the obscura group are collapsed (green triangle; melanogaster group and brown triangle: other *Drosophila* species). Posterior Bayesian Supports are only shown in the nodes with support less than 1. Duplicated genes which could not be assigned as ancestral or duplicate is marked by _1 or _2. Scale axis is in the time relative to the obscura speciation, which was set to 1.

**Figure 5.**
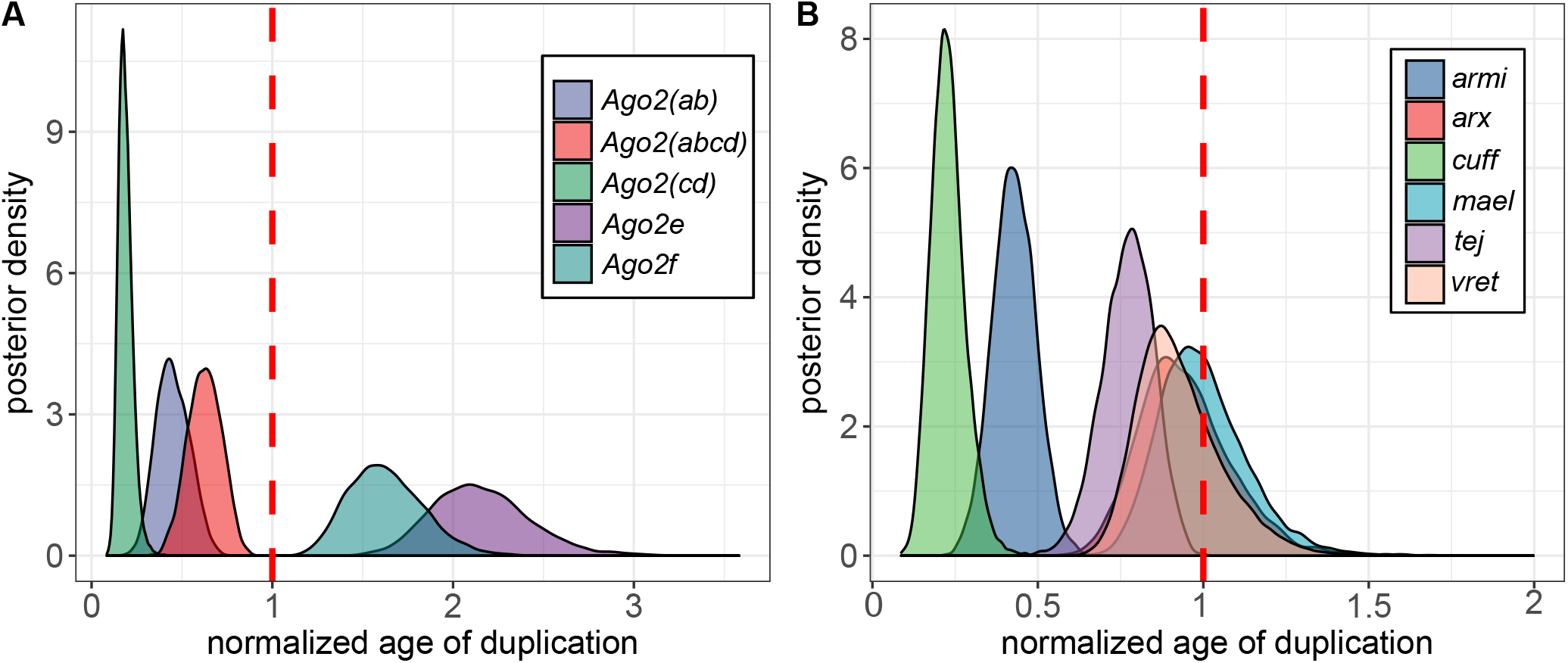
Density plot for posterior distributions of the duplication age. The MCMC posterior of the age of duplication node after 25% burn-in. (**A**) *Argonaute 2* and (**B**) RNAi accessory proteins. The broken red line denotes speciation event in *obscura* group which was normalised to be 1.

## Discussion

Although four of the six piRNA pathway duplicates did not display altered tissue specificity compared to the ancestral copy, *asterix* and *cutoff* both became significantly more male biased, as did each of the *Ago2* duplicates [4]; **Figure 1**, **Figure 2**). In each case, this was due to higher (or exclusive) expression in the testis. The duplicated genes also showed higher rates of adaptive amino acid substitution, together and individually, whereas only two (*asterix* and *armitage*) displayed evidence of positive selection when single-copy in *D. melanogaster* (**Additional file 5 Table S3**).

This new tissue specificity and the rapid evolution of duplicated copies broadly suggest that gene duplication in these pathways may be associated with functional diversification through neofunctionalization, for example by testis-specific selective pressure. Three main selective pressures seem likely candidates to have driven this process. First, given the role of *Ago2* in *Drosophila* antiviral defense [7], and the role of the piRNA pathway in antiviral defense in mosquitoes [47], it is possible that these duplications have specialized to a virus that is active in the male germline, such as *D. obscura* and *D. affinis* Sigmaviruses [48]. Second, given the role of all of these genes in the suppression of transposable elements (TEs), their evolution may have been shaped by the invasion of TEs that are more active in testis, as seen for Penelope [49] and copia [50]. Such a ‘duplication arms-race’ in response to TE invasion is thought to occur in mammals, where repeated duplications of KRAB-ZNF family are selected following the invasion of novel TEs, and subsequently provide defence [51, 52]. Alternatively, duplicates may quantitatively enhance the pre-existing response to TEs, as suggested for another rapidly-evolving piRNA-pathway component, Rhino [53].

The third, and arguably most compelling hypothesis, is that selection is mediated by conflict between meiotic drive elements and their suppressors, such as sex ratio distorting X-chromosomes [54, 55]. Most directly, meiotic drive elements are common in *Drosophila*, and RNAi-related pathways have been widely implicated in their action and suppression [15, 16, 18]. In addition, sex-chromosome drive is widespread in the obscura group: X-chromosome drive was first described in *D. obscura*, and has also been reported in *pseudoobscura*, *persimilis*, *affinis*, *azteca*, *subobscura* and *athabasca* [54] and is mediated through a testis-specific function (Y-bearing sperm have reduced function). Finally, a testis-specific class of hairpin (endo) siRNAs is required for male fertility in *D. melanogaster* [56], testes-restricted clustered miRNAs show rapid evolutionary turnover and are represented in large numbers in *pseudoobscura* [57], and suppression of sex-specific duplicates of S-Lap1 via a small-RNA mechanism has recently been implicated in the meiotic drive mechanism of *D. pseudoobscura* [58]. In this context, it is also interesting to note that *Ago2* is involved in directing heterochromatin formation in *Drosophila* dosage compensation [10–12], and that in *D. melanogaster* sex-ratio distorting *Spiroplasma* achieve male-killing through the disruption of dosage compensation (although this acts at the embryonic stage [59]). Nevertheless, in the absence of mechanistic studies, this must remain speculative, as testis is generally more permissive to gene expression and testis-specific expression may be a transient state (i.e. the “Out of Testis Hypothesis” [60]).

In this study we have built on previous work that showed *Drosophila* RNAi-pathway genes evolve rapidly and adaptively [19–22]—including multiple duplications of Ago2 in the obscura group [4]—to identify duplicates of several other RNAi-pathway genes in these species. We have shown that some of these gene duplicates have altered sex-biased expression, and that some have experienced positive selection following duplication. We suggest that this may have been driven by selection mediated through meiotic drive. Very recently, after submission of this article, improved sequencing of the *D. miranda* neo-Y and neo-X chromosomes identified a large number of previously undetected RNAi-pathway duplications (including homologs of *Argonaute 2*, *Dicer-2*, *shutdown*, *cutoff*, and *Panoramix*) within this species that also display testis-bias in their expression, further supporting a role for meiotic drive in shaping their evolution [61].

## Methods

### Sequence collation and paralog identification

The full-length sequences for 7 RNA inteference genes; *Argonaute 2* (*Ago2*), *armitage* (*armi*), *asterix* (*arx*), *cutoff* (*cuff*), *tejas (tej)*, *maelstrom* (*mael*), and *vreteno* (*vret*) from 12 obscura group species were identified using tBLASTn (BLAST+ 2.6.0) [62] with a local BLAST database (see below for the details of the construction of local genomic database). Known gene sequences from *D. pseudoobscura* and *D. melanogaster* were used as a query with a stringent e-value threshold (1e-40). Genes were inferred to have been duplicated when BLAST indicated that there were multiple full-length hits located in different genomic regions. The sequences were manually inspected, introns removed and the coding frame identified using Bioedit v 7.2 [63]. Genes in *D. pseudoobscura* were classified as ancestral or duplicate copies based on the syntenic orthology with *D. melanogaster* using Flybase Genome Browser [64]. High quality genomes are not available for other members of the obscura groups, and in those cases ancestral/derived status was assigned based on homology with *D. pseudoobscura*. To provide a comprehensive overview of the evolution of the RNAi paralogs, we included 24 *Drosophila* species outside of obscura group with assembled genomes already available in public databases. The Flybase and NCBI tblastn online portal were used to identify the target genes with queries from *D. melanogaster* or closely related species.

Five obscura group species had assembled genomes at the time of this study: *D. pseudoobscura* (assembly Dpse_3.0 [65]), *D. miranda* (assembly DroMir_2.2 [38] [66]), *D. persimilis* (assembly dper_caf1 [67]), *D. affinis* (Drosophila affinis Genome Release 1.0 [68]) and *D. lowei* (Drosophila lowei Genome Release 1.0 [68, 69]) and in these cases the genome was directly used for local BLAST database. For four species (*D. obscura*, *D. subobscura*, *D. subsilvestris*, *D. tristis*) we used de novo assembled transcriptomes based on paired RNA-seq reads data from wild-collected males [70] (Accession: PRJNA312496). Assembly was performed using Trinity [71] with ‘--trimmomatric’ and otherwise default parameters, and the assembled transcriptome was searched locally using BLAST. For three other species: *D. athabasca* [37], *D. algonquin* [37] (Accession: PRJNA274695) and *D. bifasciata* (Accession: PRJDB4817), only unassembled genomic reads were available. For these species we applied a targeted assembly approach as follows: (i) reads that had local similarity with all known duplicated RNAi proteins were identified using Diamond [72] with relaxed e-value of 1; (ii) hits from Diamond were then retained and used for assembly using Spades v3.10.1 [73]; and (iii) scaffolds produced by Spades were then used as references in local BLAST database.

### Phylogenetic analysis and the relative timing of duplications

Bayesian relaxed clock trees were used to infer the evolutionary relationship among paralogs. First, the sequences were aligned as translated nucleotide in Clustal W [74] with default parameters. Regions with ambiguous alignment were identified and removed manually by eye. A total of 7 gene trees were then inferred using Beast v1.7.0 [75]. Inference used a relaxed clock model with an uncorrelated lognormal distribution among branches, and an HKY substitution model with empirical base frequencies and rate variation among sites was modelled as a gamma distribution with four categories. The site model allowed for third codon position to have different substitution model from the other positions.

The trees were scaled by setting the time to most recent common ancestor of the *D.* obscura group to have lognormal distribution with a data-scale mean of 1, and a very small standard deviation of 0.01. This had the advantage of scaling all duplications to the same relative timescale, while allowing different genes and different paralogs to vary in their rate. To record the posterior ages of duplication, we specified the ancestral and duplicated genes as a distinct taxon set. The Monte Carlo Markov Chain analysis was run for at least 100 million states and posterior sample was recorded every 10000 states. Log files were then inspected in Tracer v1.6 [76] for parameter stationarity, and adequate sampling as indicated by an effective sample size over 200. Finally, 25% of initial trees were discarded as burn-in, and maximum clade credibility trees were summarized using Tree Annotator. Parameter MCMC files were processed using a custom R script [77] to infer the posterior distribution the age of duplication for each gene and to quantify the degree overlapping between these age distributions. To provide a reference species tree (Figure 3, in-set), we created a concatenated dataset from the ancestral copies of each duplicated gene, and inferred the gene tree in the same way.

### Differential expression analysis of the duplicated RNAi genes

For this analysis, we used obscura group transcriptome datasets available in EBI ENA (European Nucleotide Archive, http://www.ebi.ac.uk/ena) and DDBJ (DNA DataBank of Japan, http://www.ddbj.nig.ac.jp/) that included the sex and tissue annotation. The datasets comprised 163, 42 and 34 RNA-seqs datasets of *D. pseudoobscura, D. miranda* and *D. obscura* respectively; Bioproject: DRA004463, PRJEB1227 [78], PRJNA226598, PRJNA219224 [79], PRJNA326536 [45], PRJNA74723, PRJNA321079, PRJNA291085 [80], PRJNA268967 [81]. Since our main interest was the comparison expression between sex, but not its absolute expression value, we performed a simple read-counting analysis. In outline, each RNA-seq dataset was mapped to the full-length CDS using Bowtie2 v2.3.2 [82] with mode ‘--very sensitive’ and otherwise default parameters. The reads mapped to reference were counted using combination of SAMtools view flag −F 4 and SAMtools idxstats v1.4 [83]. The count data were then normalized by gene length and read depth, where it was then scaled relative to the expression of RpL32. To determine the statistical significance of difference gene expression, generalised linear mixed models were fitted using R package MCMCglmm [84] with sex as fixed effect and tissue as a random effect, and log-transformed normalised expression as the response variable. The natural logarithm transformation (log_e_) was used to reduce the skewness of the distributions. To allow for zero value for non-expressed genes, the genes with read count 0 was replaced with 1.

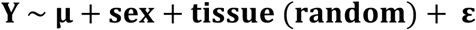

Where Y is log_e_ transformed normalized expression data (response variable), µ is mean of log_e_ transformed expression and ε is residual error. The random effects (tissue) and the residual were assumed to be distributed multivariate normal with mean 0 and uncorrelated covariance matrix MVN (0, **I**σ^2^). Sex was modelled as a factor with 2 variables (male-female) and tissue contained 13 variables of different tissue.

### Population Genetic Analysis of the RNAi Duplicated Genes

We used the McDonald-Krietman test [42] to compare the rate of adaptive evolution between ancestral and duplicate genes using polymorphism data from publicly-available sequencing datasets: Pseudobase (12 strains of *pseudoobscura*, Accession list: SRP007802 [69]) and 12 strains *D. miranda* (Bioproject: PRJNA277849 [85]).

Genomic reads for each strain were mapped to the genomic reference using Bowtie2 [82] with ‘--very-sensitive’ mode and otherwise default parameters and reads mapped to the genes of interest were extracted using SAMtools view (flag −F 4). Duplicate reads were marked using MarkDuplicates (Picard Tools [86]). To reduce the excessive variants surrounding indel, we then applied GATK IndelRealigner [86], which discards the original mapping and performs local-realignment around indel. The output was then sorted and indexed and the BAM file was used for ‘mpileup’ variant calling (SAMtools v1.4 [87]). The output VCF files were then filtered to only include SNP (GATK SelectVariants [86]), and variants that were covered by less than five reads were masked with ‘N’ (undetermined bases, --snpmask GATK v3.5 [86]). The variant files were then converted to FASTA format using GATK FastaAlternateReferenceMaker, which replaced genomic reference with variants defined in VCF files [88] and output the heterozygous calls with IUPAC ambiguous code. Finally, FastPHASE [89] was used to generate pseudo-haplotypes, although haplotype information was not utilized by the analysis.

MK tests were performed for each gene on *D. pseudoobscura-D miranda*. DNAsp v5.0 [90] was used to estimate the statistics for the MK test and Fisher’s exact test was used to calculate the statistical significance for single-gene analyses. Genes were then grouped into ancestral and duplicate genes, and a cross-gene analysis was performed using a maximum likelihood extension of the MK test [43]. Five different models were fitted that differed in the constraint of α (proportion of non-synonymous subtitutions estimated to be adaptive), and the relative support between models was compared using Akaike Weights.

## List of Abbreviations

Ago2: Agronaute
2 armi: armitage
arx: asterix
BAM: Binary Alignment Map
BLAST: Basic Alignment Search Tool
cuff: cutoff
GATK: Genome Analysis Toolkit
HKY: Hasegawa, Kishino and Yano model
mael: maelstrom
MCMC: Monte Carlo Markov Chain
MK: McDonald-Kreitman test
RNAi: RNA interference
SNP: Single Nucleotide Polymorphism
TE: Transposable Elements
tej: tejas
VCF: Variant Call Format
vret: vreteno

## Declarations

### Ethics approval and consent to participate

Not applicable

### Consent to publication

Not applicable

### Availability of data and material

Fasta alignment (both for phylogenetic and MK analysis) and raw expression data are available via Figshare (DOI: 10.6084/m9.figshare.7145720).

### Authors’ contributions

DJO and DC conceived the study and designed the analysis, DC analyzed the data and wrote the first draft of the manuscript. Both authors read and approved the final manuscript.

## Acknowledgements

We thank Billy Palmer for initial discussion on the variant calling, and Billy Palmer and Samuel Lewis for comments on an earlier version of this manuscript. We thank the many people who made their published data publicly available, and Shu Kondo for permission to use unpublished data from *D. bifasciata*.

## Additional files

### Additional file 1 Table S1

File format: xlsx

Title: **The detailed RNAi genes and its duplicate in** *D. pseudoobscura*

Description: The genomic position is based on the D. pseudoobscura assembly 3.0.

### Additional file 2 Figure S1

File format: pdf

Title: **The expression profile of RNAi across tissue**

Description: The normalized expression plotted across tissues. The error bars denote the standard error for the given tissue. Blue bar indicates male and female is indicated by red bar. The plot is shown for *D. pseudoobscura* (n=163), *D. miranda* (n=42) and *D. obscura* (n=34), respectively.

### Additional file 3 Table S2

File format: xlsx

Title: **MK test of duplicated RNAi genes in** *D. pseudoobscura-D. miranda*

Description: Ds is synonymous divergence, Dn is non-synonymous divergence, Pn is the non-synonymous polymorphisms, Ps is the synonymous polymorphisms, α represents proportion of substitutions that are adaptive, a is the absolute number of adaptive substitutions. Ln is the number of non-synonymous sites, Ls is the number of synonymous sites. Ka is the number of non-synonymous mutations per non-synonymous sites and Ks the number of synonymous mutations per synonymous sites from single randomly chosen strain for each species (Li, 1993 [91] calculated using R package seqinr). Parameter **ω_a_** is identical to Ka/Ks ratio except that the numerator only takes adaptive divergence (α * Dn)/Ln)/(Ds/Ls).

### Additional file 4 Figure S2

File format: pdf

Title: **The heat-map p-value of the difference between pairwise posterior distributions** Description: Large p-value (>0.05) indicates the overlapping distribution and the duplication time might be shared (blue box). Red boxes denote comparison with p-value < 0.05 which indicate non-overlapping posterior distribution and an asynchronous duplication event. Pink colored box indicates the marginally significant (0.01 < p-value <0.05).

### Additional file 5 Table S3

File format: multiple-tab xlsx

Title: **Additional MK test analysis**

Description: Table the results of additional MK test analysis where minor allele frequency is removed, repetition with larger dataset and results in *D. melanogaster*.

