## Supplementary figures and images for "Widespread gene duplication and adaptive evolution in the RNA interference pathways of the *Drosophila obscura* group"

### Additional file 2 Figure S1

#### *Drosophila pseudoobscura*

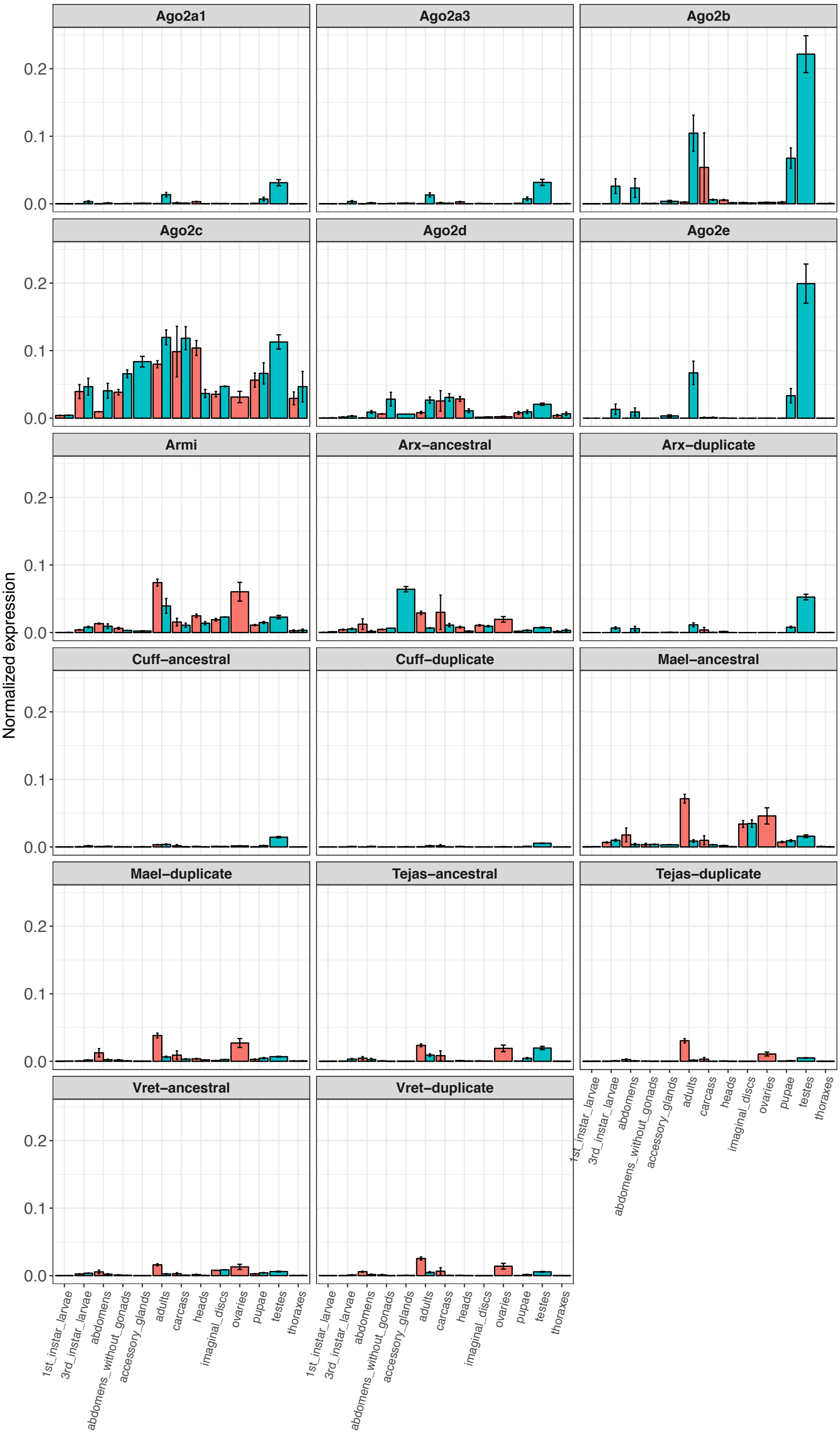

#### *Drosophila miranda*

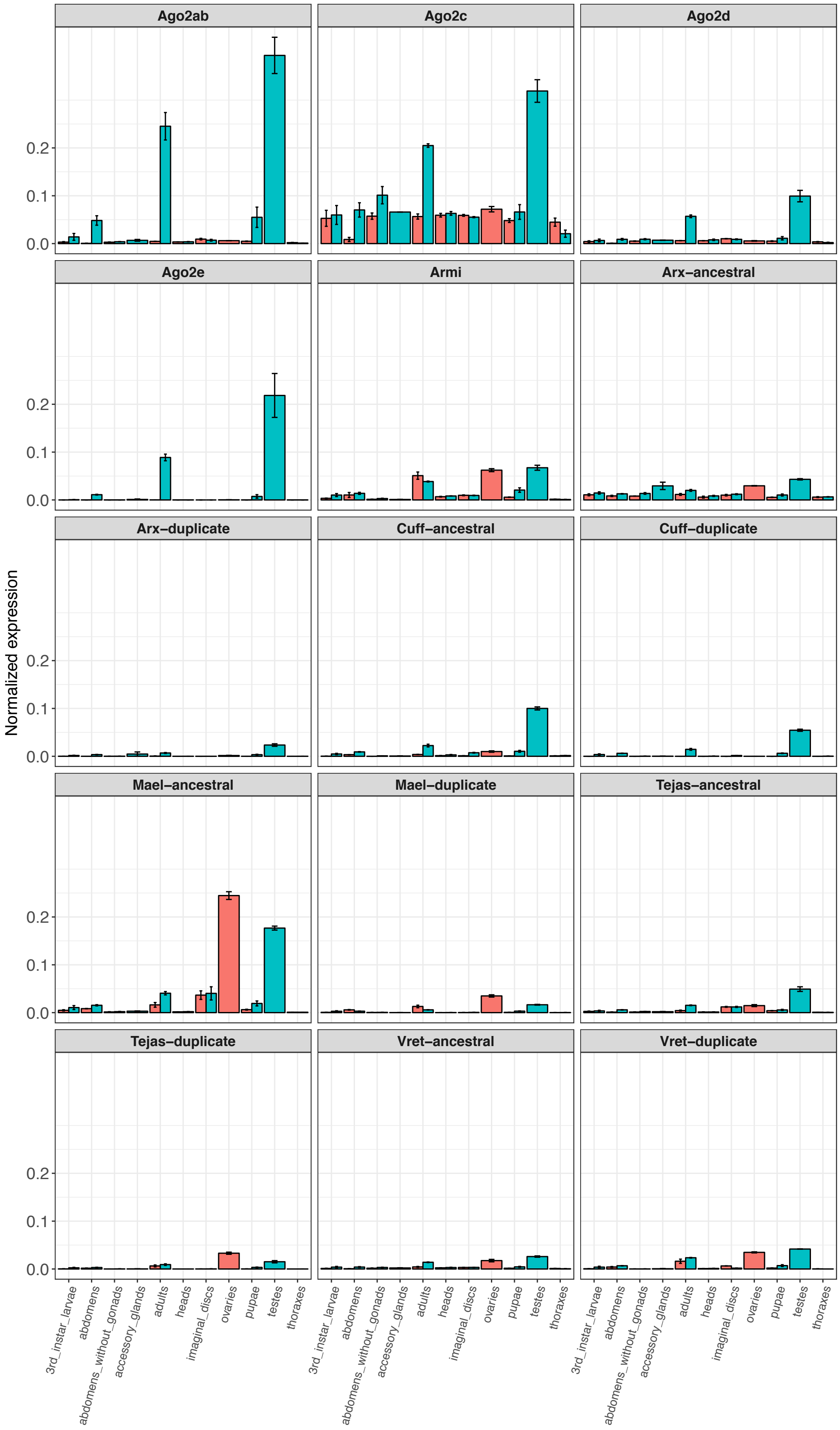

*Drosophila obscura*

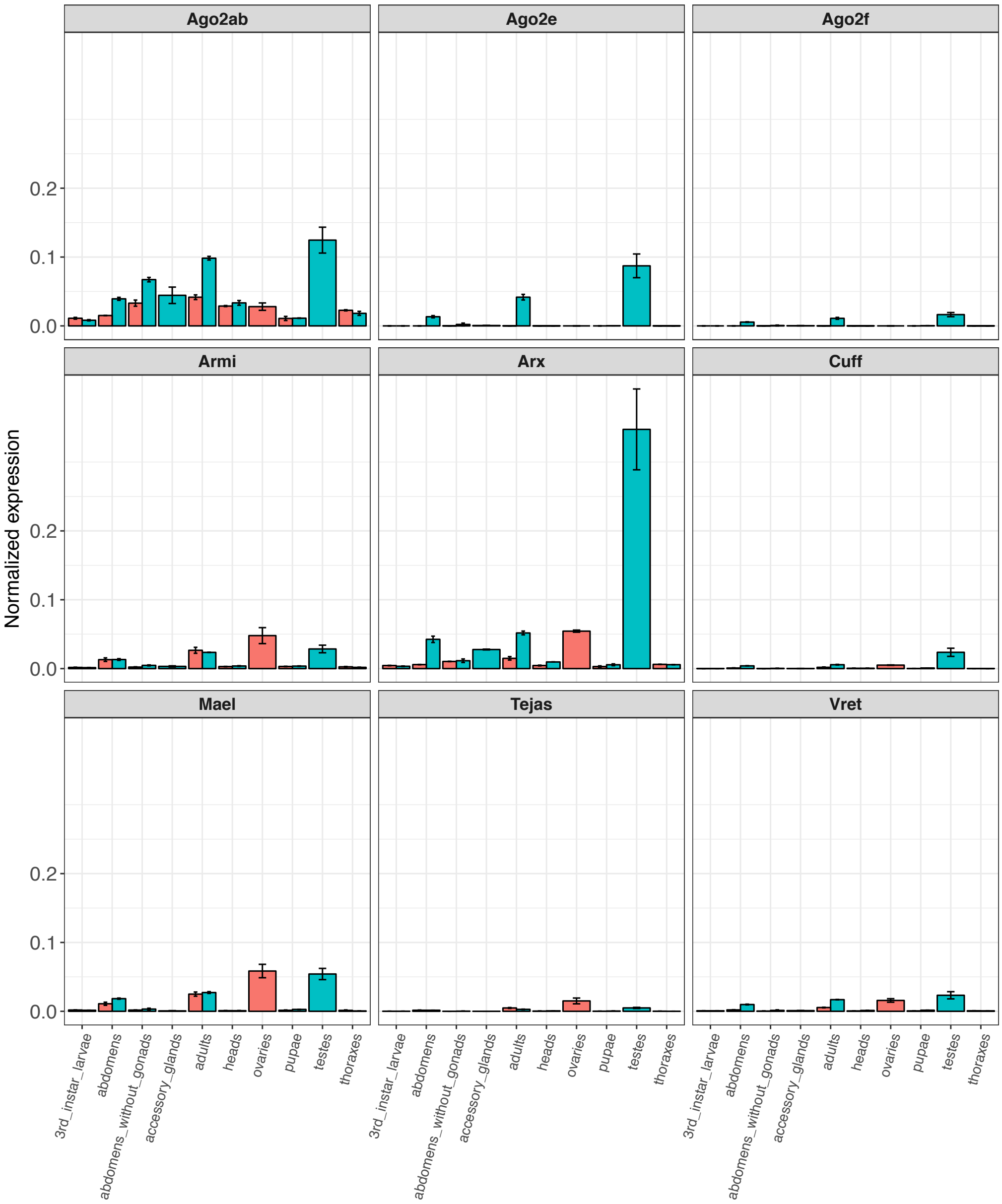

### Additional file 4 Figure S2

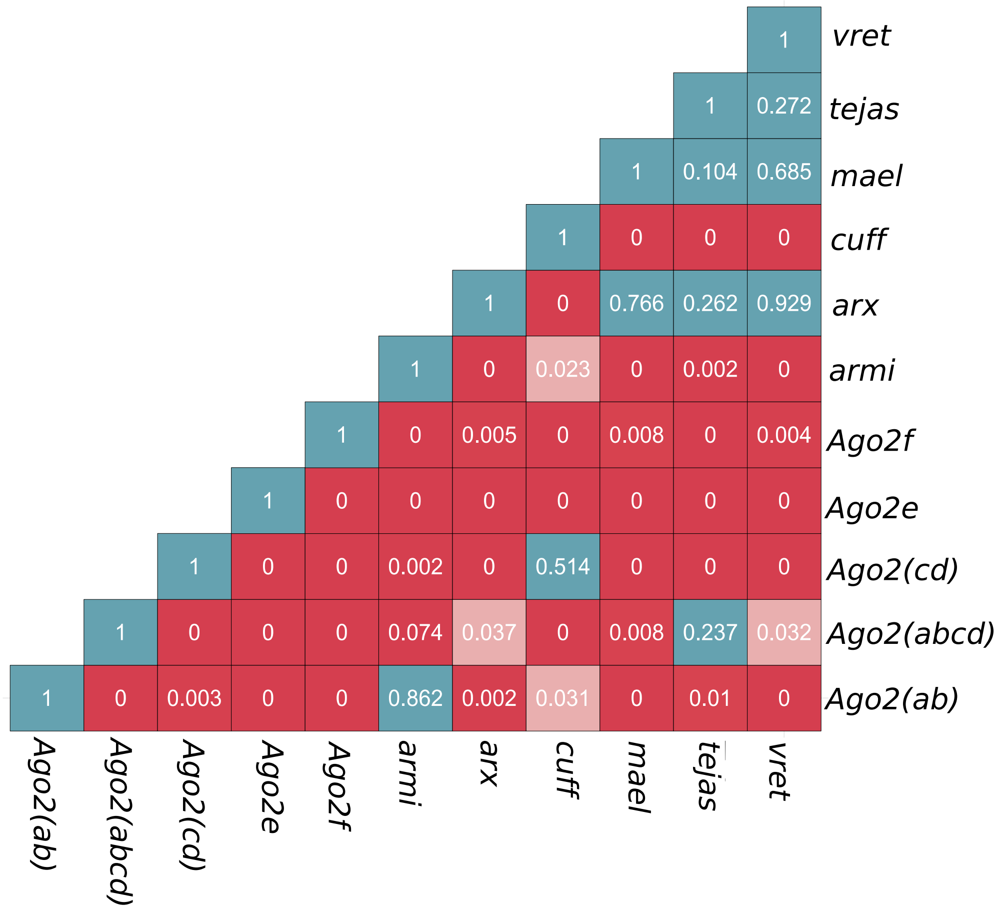
